# Impact of Cell Shape on Mitotic Spindle Positioning Forces

**DOI:** 10.1101/2023.12.15.571813

**Authors:** Jing Xie, Javad Najafi, Aude Nommick, Luc Lederer, Jeremy Salle, Serge Dmitrieff, Nicolas Minc

## Abstract

Cell geometry is a key parameter for the regulation of mitotic spindle positioning during early embryo development and tissue morphogenesis. To date, however, we still lack an understanding for how intracellular forces that position, orient or hold mitotic spindles depend on cell geometry. Here, we used *in vivo* magnetic tweezers to directly measure the forces that maintain the mitotic spindle in the center of sea urchin cells that adopt different shapes during early embryo development. We found that spindles are held by viscoelastic forces that progressively increase in amplitude as cells become more elongated during early development. By coupling direct cell shape manipulations and *in vivo* force measurements, we establish how spindle associated forces increase in dose dependence with cell shape anisotropy. Cytoplasm flow analysis and hydrodynamic simulations suggest that this geometry-dependent mechanical enhancement results from a stronger hydrodynamic coupling between the spindle and cell boundaries, which dampens cytoplasm flows and spindle mobility as cells become more elongated. These findings establish how cell shape affects spindle associated forces, and suggest a novel mechanism for shape-sensing and division positioning mediated by intracellular hydrodynamics with functional implications for early embryo morphogenesis.

## INTRODUCTION

Cell division is critical to promote the multicellularity of developing embryos and tissues. Cells usually divide at precise locations and along precise axes, a process often crucial for tissue morphogenesis and cell fate decisions (1–3). The position of the division plane is specified by the mitotic spindle, a large microtubule-based assembly that resides in the cytoplasm. Spindles can rotate or translate during mitosis to eventually adopt a final stable position that they need to maintain to specify cleavage furrow assembly (4–6). Force-generating cytoskeletal networks such as microtubule asters and associated motors or acto-myosin contractile bundles are well-established regulators of both spindle displacement and maintenance (7–12). These cytoskeleton elements connect the cell surface to spindle poles for force exertion. When cytoskeleton forces are imbalanced around cells, the spindle will translocate or rotate. When they are balanced, the spindle will hold its position and orientation. In addition to cytoskeletal force generators, bulk cytoplasm flows and viscoelastic properties, were also recently proposed to drag or hold spindles in place (13–16). Therefore, spindle positioning may result from a complex integration of different mechanical inputs.

One challenge associated with the study of mitotic spindle positioning during development, is given by the large range of cell shapes and sizes found in embryos or tissues. For instance, as early embryos develop, dividing blastomeres can vary in size over 2-3 orders of magnitude. Moving or maintaining a spindle in the center of a large versus a small cell, may for instance require very different mechanisms (17–19). Cell shape also varies extensively during development and has been shown in numerous cell types and multicellular contexts to have a major influence on spindle positioning, with spindles that commonly align along the longest cell shape axis (20–27). In general, however, how mechanical elements that regulate spindle positioning may be adapted to the size or shape of cells during development, to promote diverse division patterns remains unknown.

To address this question, we used *in vivo* magnetic tweezers to directly measure the forces that maintain mitotic spindles in the center of early dividing sea urchin blastomeres. We found that spindles are held by viscoelastic forces that progressively increase in strength from the 1- to the 2- and the 4-cell stages, as blastomeres become smaller and more anisotropic in shape. By directly manipulating cell shape, we demonstrate that cell shape anisotropy is sufficient to increase the amplitude of spindle maintaining forces. Using finite-element hydrodynamics simulations and cytoplasm flow analysis, we establish how cell shape anisotropy can enhance cytoplasm viscoelastic resisting forces on spindles, as a result of stronger hydrodynamic interactions between the spindle and cell boundaries.

## RESULTS

### Mitotic spindles are held by viscoelastic forces in the center of early blastomeres

During the early development of sea urchin embryos, as in many invertebrate and vertebrate embryos, zygotes and blastomeres divide symmetrically, by positioning mitotic spindles in the cell center (28–31). As a result of cell-cell adhesion and confinement by embryo enveloping layers, divided blastomeres also change shape significantly, and become more anisotropic, transiting from a round zygote to two hemispherical 2-cell blastomeres and to four ellipsoidal 4-cell blastomeres (29, 32) (Fig. 1A-B, Movie S1). We first sought to exploit these shape variations to test how spindle positioning forces may depend on cell geometry. To directly measure forces that hold spindles in the center of early blastomeres we applied calibrated forces on mitotic spindles along the spindle axis in zygotes, 2- and 4-cell blastomeres using *in vivo* magnetic tweezers (Fig. 1C-H, and Movie S2). These assays rely on the injection of specific magnetic beads that recruit dynein to self-target themselves to centrosomes, and form compact aggregates that stay attached to centrosomes throughout early embryo development (13, 33, 34). Upon magnetic force application on one spindle pole, spindles at all stages moved as compact objects, allowing to record a displacement-time (e.g. creep) behavior. Interestingly, these displacement-time curves exhibited similar signatures from the 1 to the 4-cell stage, indicative of similar mechanical mechanisms holding spindles in the cell center. First, the spindle moved at constant speed following a linear viscous regime, and then slowed down indicating the presence of internal elastic forces that limit spindle displacements. On longer time-scales, the spindle recovered a constant speed suggesting a progressive fluidization of elastic elements (Fig. 1I). In agreement with the existence of an elastic response, when the force was released, spindles recoiled back towards their initial position at the cell center. However, this recovery was only partial, with spindles recoiling only ∼50% of the distance moved under force, indicating a significant dissipation of stored elastic energy (Fig. S1A-B). We conclude that mitotic spindles from the 1- to the 4-cell stages are held in the cell center by viscoelastic spring-like forces.

**Figure 1.**
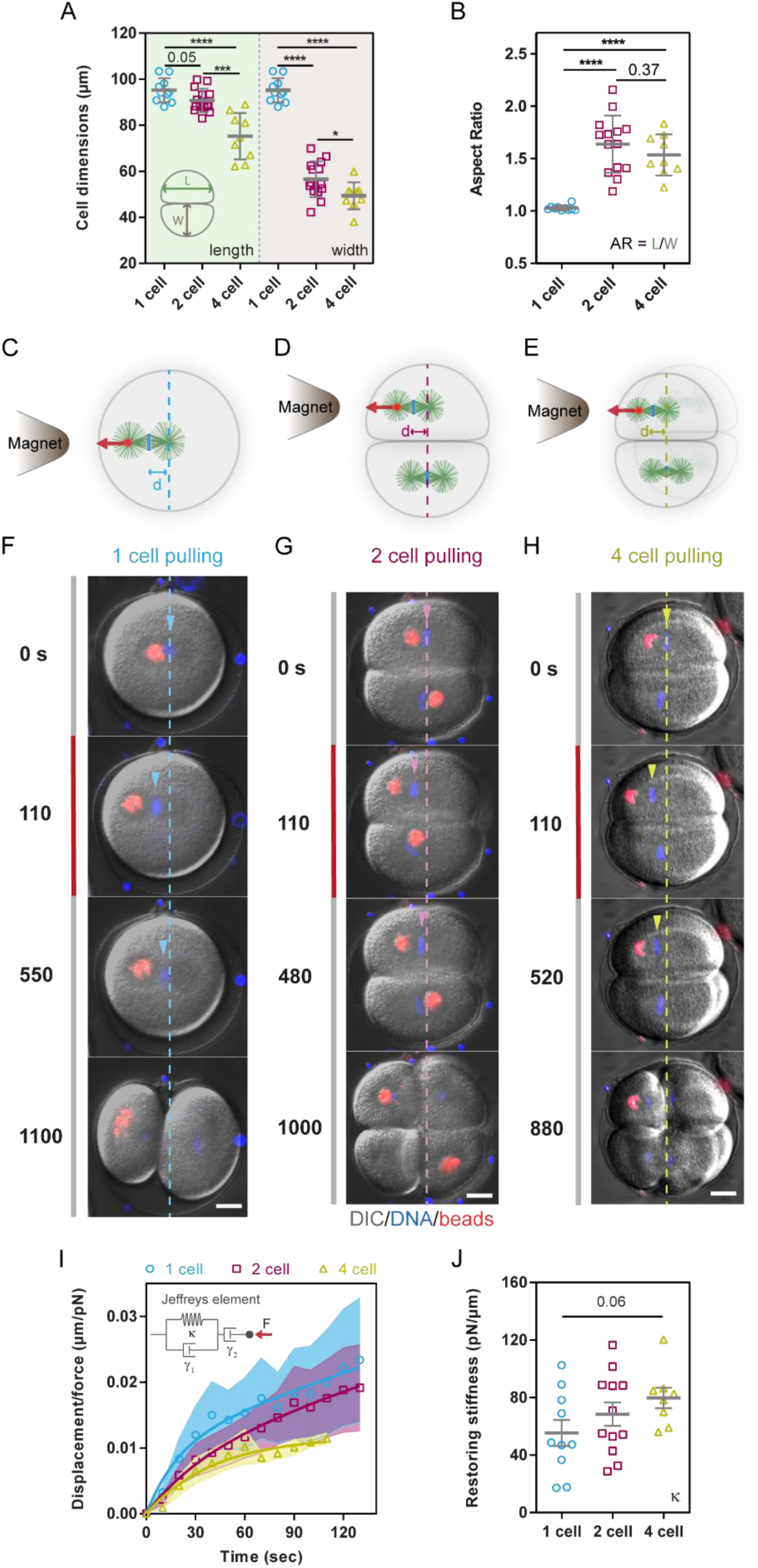
Mitotic spindles are held in the center of early blastomeres by viscoelastic forces that increase in strength with developmental stages. **(A)** Quantification of cell length and width for 1, 2 and 4-cell stage blastomeres (n= 10, 14 and 9 embryos respectively). (B) Cell shape aspect ratio of 1 ,2 and 4-cell stage blastomeres (n= 10, 14 and 9 embryos respectively). **(C-H)** Schemes and time-lapses of metaphase spindles at 1-cell stage (C and F), 2-cell stage (D and G) and 4-cell stage (E and H) with magnetic beads bound to one spindle pole, displaced by magnetic forces applied parallel to the spindle axis from the presence of a magnet tip (red line), and recoiling upon force cessation. Dotted lines correspond to the initial centered position of the spindle and arrow heads indicate the position of chromosomes. **(I)** Time evolution of the displacement measured from the initial centered position of the spindle normalized by the applied magnetic force for metaphase spindles in 1-cell, 2-cell and 4-cell embryos (n=10, 12 and 8 cells respectively). The bold lines correspond to fits of the data using a Jeffreys’ viscoelastic model (see main text and methods). Error bars are represented as shades in these curves correspond to +/- SD/2. **(J)** Restoring stiffness on mitotic spindles moved parallel to the spindle axis for the same cells and conditions as in I, computed using fits to the Jeffreys’ models (n=10, 12 and 8 cells respectively). Error bars correspond to +/- SEM. Results were compared by using a two-tailed Mann–Whitney test, and P-values are reported on the graph, or indicated as ***, P < 0.001, ****, P < 0.0001. Scale bars, 20 μm.

To compute the magnitude of these forces, we fitted rising and relaxation responses with a 3- element Jeffreys’ viscoelastic model, which provided the best fitting model for these curves (13, 35, 36) (Fig 1I). This allowed us to extract a spindle restoring stiffness, that quantifies the amplitude of elastic forces normalized by the distance the spindle moved from the cell center, as well as a spindle viscous drag, which represents the ratio between force and initial spindle velocity (7, 13). Remarkably, these analyses revealed that spindle restoring stiffness increased gradually during early development, from values of 55.34 +/- 28.8 pN/μm (+/-SD) for zygotes, to 68.41 +/- 28.21 pN/μm for 2-cell stage blastomeres, and 79.75 +/- 20.03 pN/μm for 4-cell stage blastomeres (Fig. 1I). This shows that displacing the spindle 5μm away from the cell center (few % of cell length) requires large forces that increase from ∼276 pN, ∼342 pN to ∼398 pN from the 1 to the 2- and 4-cell stages, respectively. Viscous drags also increased at subsequent stages, so that viscoelastic time-scales remained at similar values of ∼1 min independent of developmental stages, in agreement with spindle recoils occurring at the same rates (Fig. S1C-D). These results indicate a progressive rise in the forces that hold spindles in the cell center when cells become smaller and more elongated during early development.

### Cell shape anisotropy may enhance cytoplasm viscoelastic resistance on spindles

Viscoelastic forces holding mitotic spindles in the cell center were previously reported and either attributed to networks of microtubules (MTs) that reach to the cell cortex to push with length-dependent forces (7, 18), or alternatively to the mere gel-like viscoelastic properties of bulk cytoplasm that baths the spindle (13). A central difference between these two models resides in the presence (respectively, absence) of numerous MTs that reach to the cell surface (19). Therefore, in order to distinguish between these two models, we performed immunostaining to label MTs, F-actin and DNA, and used airy-scan high-resolution confocal microscopy to measure the distance between MT +TIPs and the actin-rich cortex in metaphase from the 1- to the 16-cell stage (Fig. 2A and Fig. S2A-B). The mean distance was 14.48 +/- 2.65 μm (+/-SD) in zygotes, and gradually reduced at subsequent stages to reach a distance of only 2.54 +/- 2.34 μm at the 16-cell stage (Fig. 2B and Fig. S2C). These distances represented a relatively stable fraction of ∼15% of cell length from the 1 to the 4-cell stage, but sharply transited to only ∼6 % of cell length from the 8-cell stage onward (Fig. 2C and Fig. S2D). In addition, the number of MTs reaching a distance less than 2μm of the actin-rich cortex was 0, 0.4 and 3.5 MTs/cell at the 1-, 2 and 4 cell stages, but increased to 17.88 and 24.33 MTs/cell at the 8- and 16-cell stages (Fig. S2C). These analyses suggest that spindles are uncoupled from the cell surface up to the 4-cell stage in this system, favoring a primary role for cytoplasm viscoelasticity in holding spindles in the center of early blastomeres.

**Figure 2.**
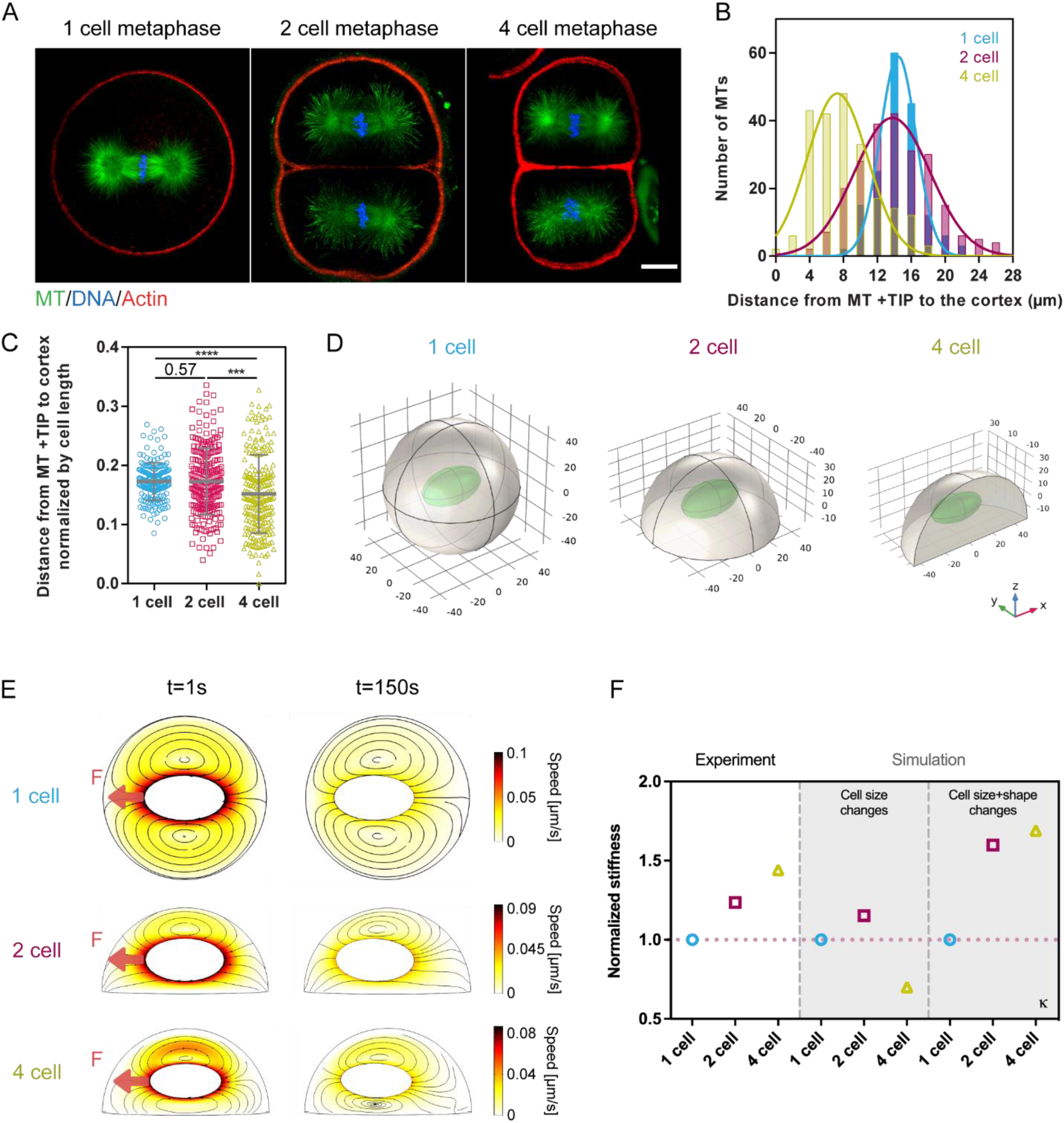
Modulations in spindle restoring stiffness with cell shape may rely on cytoplasm viscoelasticity and changes in hydrodynamic boundary conditions. **(A)** Airy-scan confocal images of sea urchin embryos at 1-cell (left), 2-cell (middle) and 4-cell stage (right) in metaphase fixed and stained for Microtubules (MTs), DNA and F-actin. **(B)** Quantification of the distance from MT+ TIPs to the actin cortex in 1-cell, 2-cell and 4-cell stage embryos at metaphase (n= 168, 229 and 220 MTs from 4 independent embryos, respectively). **(C)** Quantification of the distance from MT+ TIPs to the actin cortex normalized by cell length for the same cells and conditions as in B). **(D)** Schematic of cell and spindle geometries at different developmental stages taken to simulate the response of spindles to applied forces in a viscoelastic cytoplasm. **(E)** Stream lines and fluid flow speeds corresponding to 2 time-points (beginning and end of pulling) of hydrodynamic simulations of spindles moving under force in a viscoelastic medium confined by cell boundaries at different developmental stages. **(F)** Restoring stiffness normalized by the 1-cell stage value compared between data from experiments and different sets of simulation considering changes in cell size only or in both cell size and shape. The red dotted line corresponds to a value of 1. Results were compared by using a two-tailed Mann–Whitney test. P-values are reported on the graph, or indicated as ***, P < 0.001, ****, P < 0.0001. Scale bars, 20 μm.

To understand if and how viscoelastic forces associated to bulk cytoplasm material properties could increase in amplitude as cells become smaller and more elongated, we implemented finite-element hydrodynamic simulations (14, 36). The spindle was modeled as a rigid impermeable ellipsoidal object with dimensions taken from experimental measurements at different developmental stages (Fig. S2E). Cell shapes and sizes were obtained by cutting the 1-cell sphere into two hemispheres for the 2-cell stage and into four quarters for the 4-cell stage (Fig. 2D and Table S1). The cytoplasm was implemented as a viscoelastic fluid using an Oldroyd-B model, which is equivalent to the Jeffreys’ model used to analyze experimental curves, obeying no-slip conditions at the interface with both spindle and cell boundaries (36). To simulate magnetic pulls on spindles, a constant step force was applied to one half of the spindle using a boundary load. At the beginning of each simulation, the spindle was placed in the center of mass of the geometry, and also aligned with the longest shape axis, as in experiments. We simulated both pulling and relaxation, and fitted the displacement-time curve of the spindle, as in experiments, to compute simulated values of restoring stiffnesses and viscous drags. In addition, these simulations allowed us to predict cytoplasm flow fields and amplitudes that accompany spindle movements in the cell interior.

To isolate the sole effect of cell size that decreases during early development, we first ran simulations in spheres of reducing volumes and scaled the spindle according to experimental measurements, but kept other parameters identical (Fig. 2I and Table S1). These simulations did not predict any systematic increase in restoring stiffness or drags as observed in experiments, presumably because spindles scale with cell size during early sea urchin embryogenesis. Next, we simulated spindle response by inputting both cell shape and size variations as in experiments (Fig. 2E and Movie S3). Remarkably, these simulations predicted that changes in cell shapes, as observed in early blastomeres, can lead to a progressive increase in both spindle restoring stiffnesses and drags in close agreement with experimental behavior (Fig. 2F and Fig. S2F). These simulation results suggest that an anisotropic cell shape may be sufficient to enhance cytoplasm mechanical resistance on mitotic spindles.

### Manipulation of cell shape is sufficient to alter cytoplasm viscoelastic resistance on spindles

To test the impact of cell shape on spindle positioning forces, we first applied forces orthogonal to the spindle axis and compared the response between round isotropic zygotes and elongated 2-cell stage blastomeres. In both cells, this mode of force exertion caused spindles to translate as well as rotate to reorient towards the magnet tip (Fig. 3A-D and Movie S4). Strikingly, both translating restoring stiffness and drags were significantly increased by 168.8% and 166.1% respectively between the zygotes and the 2-cell stage blastomeres. Simulations reproduced this increase in both stiffness and drags, yielding relative values matching experimental measurements (Fig. 3E-H, Fig. S3A-S3B and Movie S5). In addition, in both simulations and experiments, in the 2-cell elongated blastomere, when the force was released, the spindle rotated back to realign along the long cell shape axis. This demonstrates that the cytoplasm can also apply restoring torques on spindles, to spring them back along the long cell shape axis (Movie S4 and S5). Therefore, spindle maintaining forces both along and orthogonal to the spindle axis become stronger with cell shape anisotropies.

**Figure 3.**
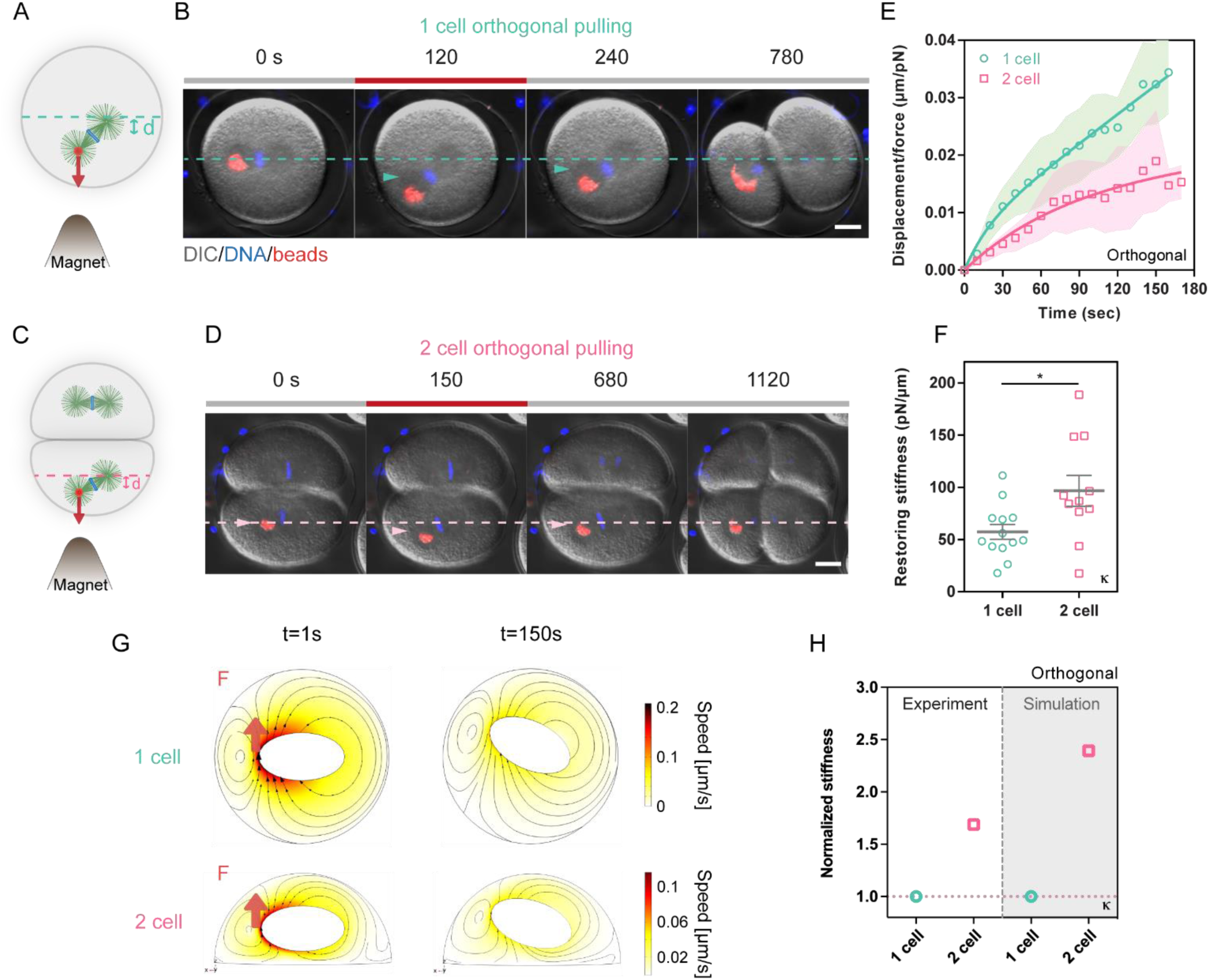
Cell shape anisotropy enhances spindle restoring stiffness orthogonal to the spindle axis. **(A-D)** Schemes and time-lapses of metaphase spindles at 1-cell stage (A and B) and 2-cell stage (C and D) with magnetic beads bound to one spindle pole, displaced by magnetic forces applied orthogonal to the spindle axis from the presence of a magnet tip (red line), and recoiling upon force cessation. Dotted lines correspond to the initial position and orientation of the spindle and arrow heads indicate the position of the pulled spindle pole. **(E)** Time evolution of the displacement measured from the initial centered position of the spindle normalized by the applied magnetic force for metaphase spindles in 1-cell and 2-cell embryos (n=14 and 10 cells respectively). The bold lines correspond to fits of the data using a Jeffreys’ viscoelastic model (see main text and methods). Error bars are represented as shades in these curves correspond to +/- SD/2. **(F)** Restoring stiffness on mitotic spindles moved orthogonal to the spindle axis for the same cells and conditions as in E, computed using fits to the Jeffreys’ models (n=13 and 11 cells respectively). Error bars correspond to +/- SEM. **(E)** Stream lines and fluid flow speeds corresponding to 2 time-points (beginning and end of pulling) of hydrodynamic simulations of spindles moving under a force orthogonal to the spindle axis in a viscoelastic medium confined by cell boundaries in rounded zygotes vs elongated 2-cell blastomeres. **(H)** Restoring stiffness normalized by the 1-cell stage value compared between data from experiments and simulations. The red dotted line corresponds to a value of 1. Results were compared by using a two-tailed Mann–Whitney test. P-values are reported on the graph, or indicated as, *, P < 0.05. Scale bars, 20 μm.

To more directly test the contribution of cell shape anisotropy to the amplitude of forces that hold spindles in the cell center, independently of developmental stages, we next designed experiments to directly manipulate cell shapes and compute forces that hold spindles. We first experimentally rounded 2-cell stage blastomeres using a transient treatment with Calcium-free sea water, that disrupts cell-cell adhesion. This caused initially elongated blastomeres to round within few minutes, without affecting cell volume, metaphase duration or spindle size (Fig. 4A-B and Fig. S4A-D). By applying magnetic pulls to mitotic spindles in these round 2-cell stage blastomeres, we found a reduction of both spindle restoring stiffness and viscous drag of 0.83X and 0.67X, respectively, as compared to those of control elongated blastomeres (Fig. 4C-E, Fig. S4E-F and Movie S6). Accordingly, hydrodynamic simulations performed using the geometry of these rounded 2-cell blastomeres also predicted a reduction in both stiffness and drag, as compared to elongated controls (Fig. 4F and Fig. S4G). These results indicate that a rounded cell geometry may alleviate cytoplasm viscoelastic resistance to facilitate spindle displacements.

**Figure 4.**
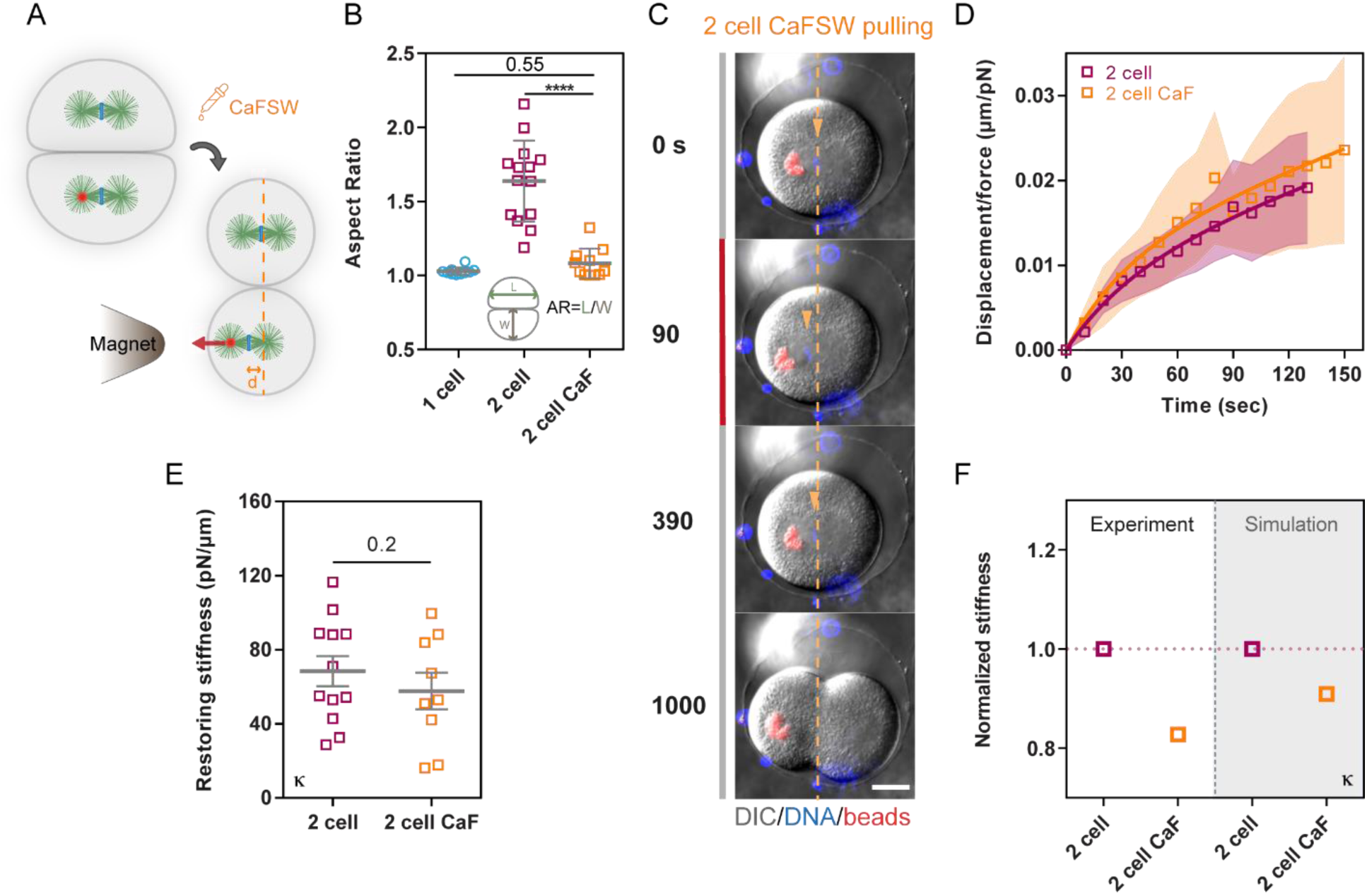
A rounded cell shape alleviates cytoplasm viscoelastic resistance on spindles. **(A)** Schemes of metaphase spindles in 2-cell stage blastomeres treated with CaFSW, to round up cell shapes with magnetic beads bound to one spindle poles. **(B)** Aspect ratios measured as indicated in the inset for cells in different conditions or stages (n=10,14 and 11 cells respectively). (C) Time-lapse of spindles in 2-cell stages blastomeres treated with CaFSW to round cell shapes, displaced by magnetic forces applied parallel to the spindle axis from the presence of a magnet tip (red line), and recoiling upon force cessation. Dotted lines correspond to the initial centered position of the spindle and arrowheads indicate the position of chromosomes. **(D)** Time evolution of the displacement of metaphase spindles at 2-cell stage normalized by the applied force for controls and cells treated with CaFSW (n=12 and 10 cells respectively). Error bars are represented as shades in these curves and correspond to +/- SD/2. **(E)** Cytoplasm restoring stiffness on mitotic spindles for the same cells and conditions as in D (n=12 and 10 cells respectively). Error bars correspond to +/- SEM. **(F)** Restoring stiffness normalized by the 2-cell control value in indicated conditions for experimental data vs simulations. Results were compared by using a two-tailed Mann–Whitney test. The corresponding P-values are indicated in the plot, or indicated as, ****, P < 0.0001. Scale bars, 20 μm. Error bars correspond to +/- SD unless otherwise indicated.

We next performed the converse experiment, and increased cell shape anisotropy to test its impact on spindle positioning forces. Starting with round zygotes, we devised U-shaped microfabricated chambers of various dimensions made in SU-8 resin that protrude above a cover-glass allowing to alter cell geometry while applying magnetic forces with the tweezer system (Fig. 5A and Movie S7). Eggs were gently pushed into chambers using a glass needle, injected with magnetic beads, and then fertilized (Fig. S5A-C). To explore dose-dependent effects, we used two sets of chambers in which cells were pushed into elongated shapes of mean aspect ratio of 1.24 and 1.49, respectively (Fig. 5A-E). In chambers, metaphase spindles moved with external magnetic forces exhibited a displacement that reflected a viscoelastic behavior similar to control, and partially relaxed back towards the cell center when the force was removed (Fig. 5F, Fig. S5E, and Movie S7). Interestingly, the measured restoring stiffness was significantly higher than in control round zygotes, increasing by 1.29X in the first set of chambers and by 1.57X in the more anisotropic chambers (Fig. 5G). Viscous drags were also increased by 1.27X and 1.22X, respectively (Fig. S5D). By performing simulations using the 3D rectangular cuboid shape of eggs pushed in the two sets of chambers we predicted an increase of restoring stiffness as well as viscous drags similar to experiments in the two sets of chambers (Fig. 5H and Fig. S5F-H). These results directly support a significant impact of cell shape anisotropy on the amplitude of viscoelastic forces that hold spindles in the cell center.

**Figure 5.**
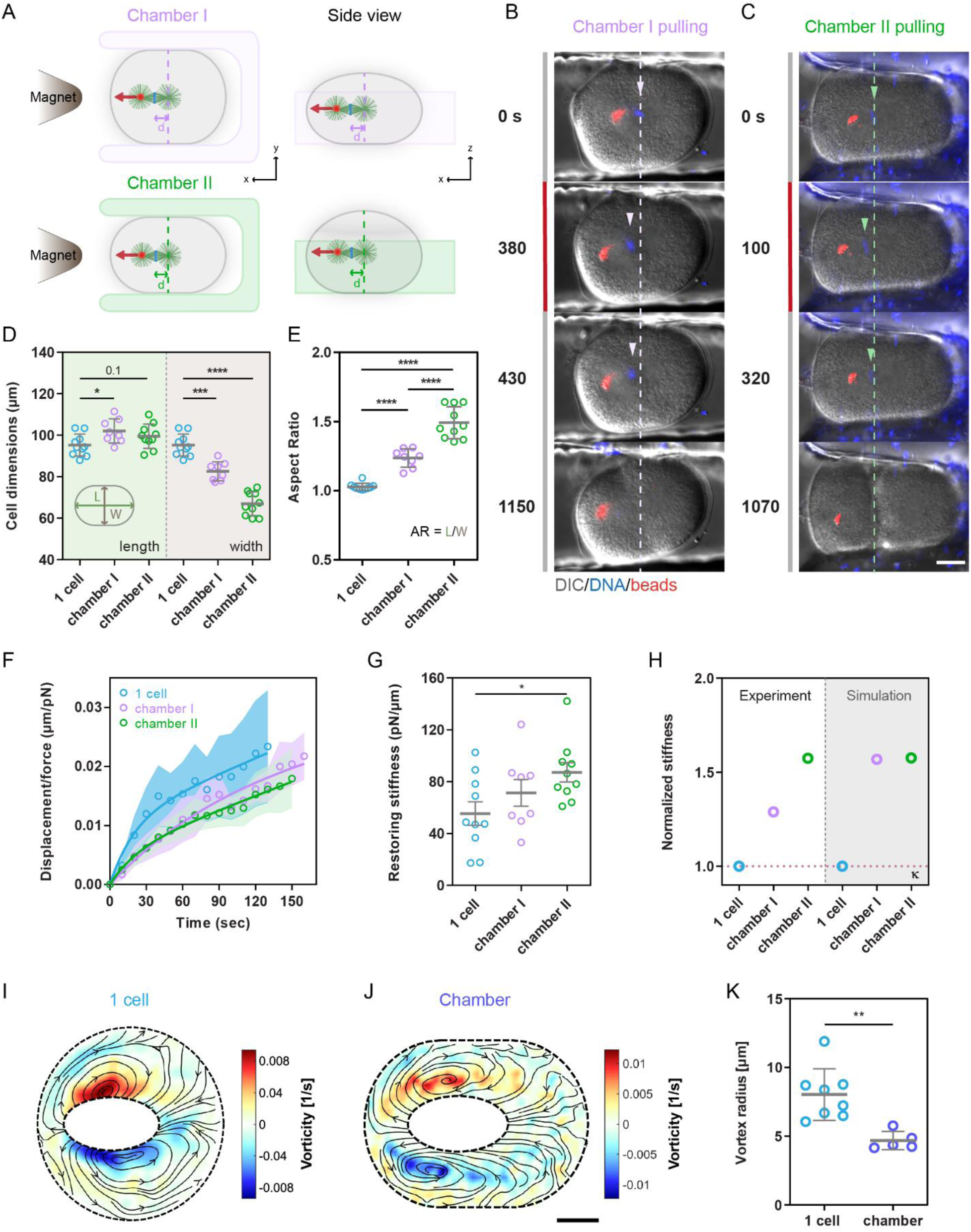
Cell shape anisotropy enhances the viscoelastic resistance of bulk cytoplasm on mitotic spindles. **(A)** Schemes representing experiments designed to change cell shapes by forcing zygotes in chambers of different geometries. (**B-C)** Time-lapses of metaphase spindles in cells shaped in chamber I (B) and chamber II (C) with magnetic beads bound to one spindle pole, displaced by magnetic forces applied parallel to the spindle axis from the presence of a magnet tip (red line), and recoiling upon force cessation. Dotted lines correspond to the initial centered position of the spindle and arrowheads indicate the positions of chromosomes. **(D-E)** Cell size and aspect ratio quantified as indicated in the inset for cells in control and in chambers (n=10, 8 and 10 cells respectively). **(F)** Time evolution of the displacement normalized by applied magnetic force of metaphase spindles at 1-cell stage in control, in chamber I and in chamber II (n=10, 8 and 10 cells respectively). Error bars are represented as shades in these curves and correspond to +/- SD/2. **(G)** Restoring stiffness of mitotic spindles for the same cells and conditions as in F (n=10, 8 and 10 cells respectively). Error bars correspond to +/- SEM. **(H)** Restoring stiffness normalized by the 1-cell value in indicated conditions for experimental data vs simulations. **(I-J)** Experimental heatmaps of cytoplasm streamlines averaged on a duration of 50 s for spindles pulled at 1-cell stage in spherical control cells (I) and in cells deformed in a chamber (J). Streamlines are superimposed onto a color map of local flow vorticities, ω. **(K)** Quantification of flow vortex radius for normal cells and cells deformed in chambers (n=8 and 5 cells respectively). Results were compared by using a two-tailed Mann–Whitney test. The corresponding P-values are indicated in the plot, or indicated as *, P<0.05, ***, P < 0.001, ****, P < 0.0001. Scale bars, 20 μm.

### Cell shape anisotropy dampens cytoplasm viscoelastic flows around moving spindles

Inspection of cytoplasm viscoelastic flows generated along moving and recoiling spindles in simulations, revealed the presence of specific flow streams and recirculation which appeared to be influenced by different boundary conditions in varying cell shapes (Movies S3 and S5). In particular, the presence of boundaries appeared to limit cytoplasm flow speed in between spindles and cell cortices, providing a potential hydrodynamic mechanism for how cell shape could impact spindle mobility. To test if flow amplitude and arrangements could be impacted by cell shape in experiments, we used Particle Image Velocimetry of large granules in cells, to map flows created along spindle pulls in round and shape manipulated zygotes (13, 36). In both control round zygotes and deformed ones in microchambers, flows exhibited two large vortices on the two sides of moving spindles, indicative of large cytoplasm fluid shear in between the cell and spindle boundaries (Fig. 5I-J). Interestingly, the size of vortices, measured as the width of the gradient in flow velocities, was much smaller in eggs deformed in chambers as compared to control round eggs, indicating a much higher shear when cell shapes become more anisotropic (Fig. 5K and S6A). Accordingly, by simulating fluid flows caused by spindle movements in diverse cell geometries, we predicted a dose-dependent decrease in the size of vortices and thus an increase in fluid shear with cell shape anisotropy, suggesting that narrower cell cross-section can dampen fluid flows to increase cytoplasm viscoelastic resistance on spindles (Fig. S6B-E). Finally, to generalize the influence of cell geometry on cytoplasm viscoelastic resistance on spindles, we ran a series of simulations in which we systematically altered cell geometry and computed spindle restoring stiffness and drags. This showed that the area, but also the aspect ratio of the cell cross-section orthogonal to the spindle axis, may be important parameters to predict viscoelastic enhancement, with narrower areas and higher aspect ratio that yielded to increased viscoelastic resistance (Fig. S6F-H). We conclude that cell shape can impact spindle positioning forces, through hydrodynamic coupling between the spindle and the cell surface, that become more prominent as cells become more elongated and narrower.

## DISCUSSION

The forces that position and orient mitotic spindles are fundamental regulators of division plane positioning and consequent tissue morphogenesis. However, these forces have only been rarely measured, given the technical difficulties of directly attaching force probes on spindle poles in intact live cells (7, 13, 37). Here, we employed a magnetic tweezer set up to directly measure these forces in intact embryos at multiple developmental stages. We found that spindles are held in the center of early blastomeres by large viscoelastic restoring forces, that reach several hundreds of picoNewtons when spindles are offset by only few microns. These forces oppose and limit spindle displacement and partially restore its initial position or orientation upon perturbation. Analysis of astral MTs length suggests that spindles lack direct contact with the surface up to the 4-cell stage. Together with signatures of spindle displacement under force, and hydrodynamic simulations predictions, these data suggest that most of these restoring forces may be attributed to bulk cytoplasm viscoelasticity that act as a soft gel that holds spindles in the cell center, and spring it back when displaced (13). Importantly, we find that these recentering forces increase in magnitude with developmental stages, as a consequence of a progressive increase in shape anisotropy in early blastomeres. Such increase may have functional implication for spindle stability during development. It may for instance allow to maintain a relative spindle centering precision mostly constant as cells become smaller (7, 18), or limit the impact of large surface stresses associated to acto-myosin contractility or cell-cell re-adhesion inherent to cell division and shape changes on spindle positioning (15, 16, 38).

Using direct cell shape manipulations, we demonstrate that an increase in shape anisotropy results in a dose dependence increase in the forces that maintain the spindle in the cell center. Simulations suggest that when cells become more elongated, their cross-section becomes narrower, limiting space for the cytoplasm fluid to flow around spindles, thereby enhancing viscoelastic resistance, akin to a piston effect (36, 39, 40). Therefore, an important output of this study is to put forward a “cell shape sensing” mechanism based on the influence of cellular boundaries on cytoplasm hydrodynamics with direct consequences on the mechanics of organelle positioning. As cell shape anisotropy typically increases during early cleavage in many embryos (32), we anticipate this regime to be highly relevant to the mechanics of spindle positioning in early dividing blastomeres. In addition, such mechanisms could also impact organelle positioning, in many other cell types, such as in elongated columnar epithelial cells that line embryo surfaces later in development (41).

This hydrodynamic shape sensing mechanism may have important generic implications for spindle positioning. First, as demonstrated in our assay in which we force cells to adopt a rounded shape (Fig. 4), the net mechanical forces that hold spindles in the cell center are alleviated in rounded cells, suggesting that a rounded geometry may effectively facilitate spindle translation or rotation. As many cell types round up during mitosis (24, 42), such effect could ease spindle rotation or asymmetric displacement needed for oriented or asymmetric divisions. Second, our measurements provide direct quantitative support for an increase in spindle centering forces as cells become more elongated. Such finding may explain the dose-dependence influence of cell geometry over polarity in orienting spindles described in multiple systems (25, 29, 43). Third, enhancing cytoplasm viscoelastic resistance in elongated cells, may help hold spindles in the cell center, when the cell’s longest dimensions become too large for cytoskeletal based system to interact with the cell surface along this axis (44). Lastly, we show that cytoplasm viscoelastic flows can apply restoring torques on spindles to realign them with the long cell shape axis, providing an important mechanism for long axis division rules (e.g. Hertwig’s rules (23)) in acentrosomal spindles that lack MT asters to sense cell geometry (45, 46).

Finally, by tracking recirculating cytoplasm flows generated along spindle displacements and recoils, we directly evidence how cellular boundaries impact the amplitude and geometries of flows in living cells. Cytoplasm flows have been mapped in organisms ranging from algae to plant and animal embryos and shown to have fundamental implications for cytoplasm homogenization, organelle transport or multicellular morphogenesis (15, 47–50). In this context, our findings document the important contribution of cell shape and size to the amplitude of streaming flows, and suggest that under a given stress, flows may be significantly dampened by the confinement imposed by specific cellular geometries. Future work addressing in quantitative terms how cytoplasm fluid mechanics intersects with cell geometry will bring important insights into mechanisms of organelle positioning and tissue morphogenesis.

## ACKNOWLEDGMENT

We thank all members of the Minc team for discussion and technical help. J.X. acknowledges the “École Doctorale FIRE - Program Bettencourt”, a fellowship from the Chinese Scholarship Council (201708070046) and from the labex “Who Am I”. J.N. is supported by a Post-doc fellowship from the ARC foundation (PDF20191209818) and is an EMBO Non-Stipendiary Fellow (ALTF 881-2019). A.N. is supported by a Post-doc fellowship from the ARC foundation (PDF2021070004082). This work was supported by the Centre National de la Recherche Scientifique (CNRS), the Université de Paris, and grants from La Ligue Contre le Cancer (EL2021.LNCC/ NiM), the Agence Nationale pour la Recherche (ANR, “TiMecaDev”), the Fondation Bettencourt Schueller (“Coup d’élan”), a scientific subvention from the foundation “Simone and Cino Del Duca” and the European Research Council (ERC CoG “Forcaster” no. 647073 and ERC PoC “MagMech” No. 101069173) to N.M.

## AUTHOR CONTRIBUTIONS

Conceptualization, N.M., J.X. and J.N.; Methodology, N.M., S.D., J.S., A.N., L.L, J.N. and J.X. Writing –Original Draft, N.M., J.X. and J.N. Draft Editing. N.M., S.D., J.S., A.N., L.L, J.N. and J.X.

## MATERIAL AND METHODS

### Sea urchin gametes

Purple sea urchins (*Paracentrotus lividus*) were obtained from the Roscoff Marine station (France) and maintained at 16°C in artificial sea water (ASW; Reef Crystals, Instant Ocean). Gametes were collected by intracoelomic injection of 0.5 M KCl. Sperm was collected undiluted, kept at 4°C and used within one week. Eggs were rinsed twice with ASW, kept at 16 °C, and used on the day of collection. Unfertilized eggs were transferred on protamine-coated glass-bottom dishes (MatTek Corporation) after removing the jelly coat through an 80-μm Nitex mesh (Genesee Scientific), injected and fertilized under the microscope.

### Immunostaining

Immunostaining was performed using procedures described previously (51). First, to remove the fertilization envelope and facilitate Antibody penetration, eggs are fertilized in a fresh solution of p-aminobenzoic acid (PABA) in ASW for 3 minutes, washed 3 times in ASW, and incubated at 16°C until the desired stage. Then, samples were fixed for 70 min in 100 mM Hepes, pH 6.9, 50 mM EGTA, 10 mM MgSO4, 2% formaldehyde, 0.2% glutaraldehyde, 0.2% Triton X-100, and 400 mM glucose. To reduce autofluorescence, eggs were then rinsed 3 times in PBS and placed in 0.1% NaBH4 in PBS freshly prepared, for 30 min. Eggs were rinsed with PBS and PBT (PBS + 0.1% TritonX) and blocked in PBT supplemented with 5% goat serum and 0.1% bovine serum albumin (BSA) for 30 min. Samples were rinsed with PBT before adding primary antibodies. For MT staining, cells were incubated for 48 h at 4°C with either a mouse anti-α-tubulin (DM1A; Sigma-Aldrich) primary antibody at 1:5000 in PBT, or a YL1/2 rat anti-tubulin (Abcam) primary antibody at 1:1000 in PBT, rinsed 3 times in PBT and incubated for 4 h at room temperature with anti-mouse or anti-rat secondary antibody coupled to Dylight 488 (Thermo Fisher Scientific), or an secondary alpaca nanobody anti-mouse 488 SMS1AF488-1-100 (Thermo Fisher Scientific) at 1:1000 in PBT. DNA was stained with 10 μg.ml^−1^ Hoechst 33342 (Sigma-Aldrich) in PBT during the same step as for the secondary antibody. For actin staining, samples were incubated for 1h in a solution of Rhodamine Phalloidin at 4U/ml in PBT. Eggs were washed three times in PBT then twice in PBS, transferred in 50% glycerol in PBS, and finally transferred into mounting medium (90% glycerol and 0.5% N-propyl gallate PBS). For staining from 1-cell stage to 16-cell stage, samples were fixed at different times after fertilization, with a time interval of around one hour. Fixation times for metaphase cells at different stages were performed at 45-55 min post-fertilization for the 1-cell stage, 1h45-1h55min for the 2-cell stage, 2h45-2h55min for the 4-cell stage, 3h45-3h55min for the 8-cell stage, and 4h45-4h55min for the 16-cell stage.

### Manipulating cell shape with CaFSW

To manipulate cell shape at the 2-cell stage from a hemisphere to a sphere, magnetic bead-injected eggs were first fertilized and incubated in para-aminobenzoic acid (PABA) for 2 min to soften the fertilization envelope, and PABA was then washed out by rinsing in ASW. The softened fertilization envelop was cut to open a hole using a glass capillary. After the first cell division, interphase cells were incubated in Hyaline Extract Medium (HEM) for 2 minutes, then rinsed with calcium free sea water (CaFSW) to disassemble cell-cell junctions (protocol adapted from (52)). Cells became round and were able to separate from their sister cells after pushing a cell out of the fertilization envelope through the opening. The remaining cell inside the FE containing magnetic beads was imaged and its spindle was moved with magnetic tweezers at 2-cell metaphase. After CaFSW treatment, cells formed normal spindles and underwent normal cytokinesis with spindle length and metaphase duration similar to controls (Fig. S4).

### Fabrication of microchambers

Resin-made microchambers were produced using standard photolithography on a coverglass. A large glass coverslip (50 * 75 mm) was first cleaned with acetone, isopropanol and DI water, and then dried with compressed air, and heated at 95°C on a hotplate for 2 minutes to evaporate any remaining chemicals. After cleaning, a first layer of Omnicoat (MicroChem) was deposited with a spincoater at 4000 rpm, and then dried for 2’ at 95°C. A layer of SU8-2050 resin (MicroChem) was then deposited with a spincoater at 2500 rpm and baked for 3’ at 65°C and 9’ at 95°C. The coverslip was then exposed to UV light for 14 seconds to polymerize the resin through a photomask with different U-shape designs. The coverslip was baked again, and developed in SU8 developer solution for 5 minutes and rinsed extensively with isopropanol. Finally, the coverslip was baked at 120°C for 10 minutes and allowed to cool gradually to room temperature. A mini-wall made of Bondic UV glue was assembled around the coverslip to form a reservoir allowing to place liquid onto the chambers. Finally, the chambers were rinsed in PBS for more than 48 h to remove any soluble chemicals from the microfabrication process.

### Manipulating cell shape with microchambers

To manipulate the shape of zygotes, from a sphere to a rectangle, the resin U-chambers were first plasma-cleaned and coated with 1% protamine. Using a microforge (Narishige MF2), a non-siliconized glass needle was cut and melted to obtain a smooth tip with an opening of 20∼30 μm, then bent at 30° and installed on a secondary hydraulic micromanipulator (Narishige NAI-30). The needle was connected to a hydraulic pressure controller (Eppendorf, CellTram) and used as holding pipette. Sea urchin eggs were placed on the chamber using the holding pipette, and gently pushed to fill the chamber shape. The eggs were pushed inside the chamber and injected with magnetic beads. After fertilization, metaphase zygotes with beads attached to one spindle pole were selected for force application using magnetic tweezers. Cells inside the chambers formed normal spindles and underwent normal cytokinesis but with a slightly longer metaphase duration than controls (Fig. S5).

### Magnetic force application

Magnetic tweezers were implemented as described previously(13, 33, 34). The magnet probe used for force applications *in vivo* was built from three rod-shaped strong neodymium magnets (diameter 4 mm; height 10 mm; S-04-10-AN; Supermagnet) prolonged by a sharpened steel piece with a tip radius of ∼50 μm to create a magnetic gradient. The surface of the steel tip was electro-coated with gold to prevent oxidation. The probe was controlled with a micromanipulator (Injectman 4, Eppendorf), and mounted on an inverted epifluorescence microscope.

Super-paramagnetic particles (diameter 800nm; NanoLink; Solulink) with spontaneous MT minus end–directed motion were used to apply magnetic forces on the spindles for early blastomeres or cells shaped with CaFSW or microchambers (13, 33, 34). To prepare beads for injection, a solution of 10 μl of undiluted streptavidin-beads was first washed in 100 μL of washing solution (1 M NaCl with 1% Tween-20), and sonicated for 5 min. The beads were then rinsed in 100 μl PBS, incubated 15 min in 100 μl 2 μg/ml Atto565-biotin (Sigma-Aldrich), rinsed again in 100 μl PBS, and finally re-suspended in 20 μl PBS and kept at 4°C until use. Unfertilized eggs were placed on a protamine-coated glass bottom dish or inside a protamine-coated microchamber. The bead solution was injected using a micro-injection system (FemtoJet 4, Eppendorf) and a micro-manipulator (Injectman 4, Eppendorf). Injection needles were prepared from siliconized (Sigmacote) borosilicate glass capillaries (TW100-4) of 1 mm in diameter. Glass capillaries were pulled using a needle puller (P-1000, Sutter Instrument) and ground with a 40° angle on a diamond micro-grinder (EG-40, Narishige) to obtain a 10 μm aperture. Injection needles were back-loaded with 2 μl of bead solution before each experiment, and were not re-used. After fertilization, beads were spontaneously transported along MTs and formed a large aggregate at the center of the aster. This aggregate stayed stably at the centrosome and occasionally split at centrosome duplication. Thus, at metaphase, beads sometimes ended up on one or the two spindle poles. The targeting of beads towards the centrosomes presumably occurred in a dynein- and microtubule-dependent manner (13).

To image spindle displacement under force, unfertilized eggs stuck on protamine-coated glass dishes or inside a protamine-coated U chamber were incubated in 10 μg/ml Hoechst (Sigma-Aldrich) before injection and fertilization. At metaphase, chromosomes marked with Hoechst, lined up along the metaphase plate, and the spindle was easily visible as a dumbbell-shaped smooth region in DIC (Differential Interference Contrast), presumably because of its association with the ER and yolk granules exclusion. These DIC images allowed to select spindles that were planar and to define the long and short-axis directions for force applications. The magnet was then rapidly moved close to the eggs and held at a fixed position along the spindle axis (see movie S2). The end of metaphase was captured as the first time-point when chromosomes separated.

### Imaging

Time-lapses of spindles and 1μm magnetic beads moving under magnetic force were recorded on two inverted microscope set-ups equipped with a micromanipulator for magnetic tweezers, at a stabilized room temperature (18–20°C). The first set-up was an inverted epifluorescence microscope (TI-Eclipse, Nikon) combined with a complementary metal–oxide–semiconductor (CMOS) camera (Hamamatsu), using a 20X dry objective (Apo, NA 0.75, Nikon) and a 1.5X magnifier, yielding a pixel size of 0.216 μm. The second one was a Leica DMI6000 B microscope equipped with an A-Plan 40x/0.65 PH2 objective yielding a pixel size of 0.229 μm, equipped with an ORCA-Flash4.0LT Hamamatsu camera. Both microscopes were operated with Micro-Manager (Open Imaging).

Imaging of immunostained cells to visualize MTs or F-actin was performed on a confocal microscope (Zeiss, LSM980) coupled to an Airyscan 2 module in confocal mode with a 63X water immersion objective (NA, 1.4; C-Apo; Zeiss).

### Magnetic force calibration

Magnetic forces were calibrated *in vitro* following procedures described previously (13, 33, 34). The magnetic force field created by each magnet tip used was first characterized by pulling 2.8 μm mono-dispersed magnetic beads (Dynal) in a viscous test fluid (80% glycerol, viscosity 8.0×10−2 Pa sec at 22°C) along the principal axis of the magnet tip. Small motion of the fluid was subtracted by tracking 4 μm non-magnetic tracer fluorescent in the same solution. The speed of a magnetic bead V was computed as a function of the distance to the magnet, representing the decay function of the magnetic force, and fitted using a double exponential function.

To compute the dependence of the force on aggregate size, bead aggregates from the same beads as those used *in vivo* in sizes ranging from 2 to 8 μm, similar to that observed in cells, were pulled in the same fluid as above. The speed V_a_ was measured and translated into a force using Stokes’ law F=6πηRV_a_, where η was the viscosity of the test fluid, R the aggregate effective radius defined using the longest length L_1_ and the length perpendicular to the longest axis, L_2_, as 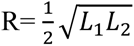. The force–size relationship at a fixed distance from the magnet was well represented and fitted by a cubic function. These speed–distance and force–size relationships were combined to compute the magnetic forces applied to spindles inside cells as a function of time, from the size of aggregates at spindle poles and their distance to magnet tips.

### Analysis of spindle position

Spindle displacement time-lapses were rotated to align the initial spindle axis to the horizontal X-axis. Magnet tip position was recorded in DIC and the position of bead aggregates was tracked from their fluorescence signal. Spindle position was processed manually in Fiji by tracking the centers of two smooth disks, which correspond to spindle poles in the DIC channel. This allowed to compute spindle length, center as well as angles with the magnetic force axis. Spindle displacements and magnetic forces were then projected along the horizontal X-axis. Chromosome plate position was also tracked but we often noticed some small delay in the motion of chromosomes as compared to that of spindle poles, presumably caused by some time-scales associated with internal elastic structures that hold chromosomes. In some experiments in which cells are treated with CaFSW, the cell moved during the pull, so the spindle position was corrected according to the displacement of the cell.

### Measuring MT distance to the cortex

Airyscan confocal images stained for MTs and F-actin from 1-cell to the 16-cell stages were projected onto a mid-section 2 μm thick, and planar spindles were selected. The horizontal and vertical distances between MT + TIPs to the actin cortex were measured, by tracing lines along each MTs to the first border of the F-actin signal using Fiji. Normalization of the distance between MTs and cortex was computed using the cell length along the spindle axis.

### Viscoelastic parameter calculation

Displacements of spindle and 1μm bead were fitted with a Jeffreys’ model using a custom code written in MATLAB (Mathworks) to compute viscoelastic parameters. For the rising phase, the spindle position or 1 μm bead position was fitted using:

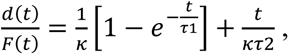

where *d* is the displacement along the X-axis, and F is the magnetic force. These fits allowed to compute the restoring stiffness, κ, and the viscoelastic time-scales, τ_1_ and τ_2_ corresponding to the Jeffreys’ viscoelastic timescales, allowing to compute viscous drags on spindles as *γ* = κτ.

Spindle relaxation dynamics was fitted using:

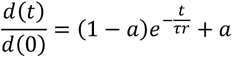

where t = 0 corresponds to the time of the end of force application, to compute the relaxation offset *a* and the relaxation time-scale τ_r_.

Fit of the data were obtained for each single force experiment, using Nonlinear least squares method in MATLAB curve fitting. Various combinations of starting points of fit parameters were tried and chosen in a way that maximum R-squared and tighter confidence interval were achieved. Curves were also visually inspected to ensure that they passed through more points and were not biased with respect to the data points.

### PIV analysis and shear measurement

The recorded image sequences of spindle pulling in DIC were analyzed using the particle image velocimetry PIVlab tool in MATLAB (Mathwork) (53). The exterior of the egg was masked to be excluded from the analysis. Contrast limited adaptive histogram equalization (CLAHE) and two-dimensional Wiener filter with accordingly windows of 20 and 3 pixels widths were applied on the images in the pre-processing steps for denoising. Image sequences were analyzed in the Fourier space by three interrogation windows with 64, 32, and 16 pixels widths and 50% overlapped area. The spline method was used for the window deformation, and subpixel resolution was obtained by two-dimensional Gaussian fits. The distribution of the velocity components of all vectors for each set was visually inspected and restricted to remove the outliers in the post-processing stage. Moreover, two other filters based on the standard deviation and local median of velocity vectors were applied to validate the vector fields. The output vector fields after smoothing were used for the analysis. The vector fields were temporally averaged over the first five frames of the pulling phase before heat map plotting and vortex analysis.

To estimate vortex size, the component of the vector field along the pulling axis x was spatially averaged over half of the egg close to the magnet after the temporal average of the velocity field. Two Gaussian functions were fitted to the side peaks of the velocity profile. These peaks started from the middle of the vortices around the spindle and extended up to the cell surface where the flow was affected by the boundary confinement. The size of vortices was calculated as the width of the Gaussian function and averaged over both sides for each sample (Fig. S6A).

### Statistical analysis

All experiments presented in this manuscript were repeated at least twice and quantified in a number of cells or events detailed in each figure legend. Statistical and correlation analyses were carried out using Prism 6 (GraphPad Software, La Jolla, CA). Statistically significant differences and tests used depended on whether experiments were paired or not and are reported in figure legends.

### COMSOL 3D simulation

The spindle pulling experiments at various cell division stages were modeled using finite element software COMSOL as described in previous studies (14, 36). The viscoelastic parameters of the cytoplasm γ_1_, γ_2_, and κ at different cell stages were measured by fitting a Jeffreys model to the creep curve of magnetic beads with a radius r=0.5 μm. These values were converted to the equivalent values in the Oldroyd-B model and inserted into the simulations as η_1_, η_2_, and τ_2_, where η_1, 2_ is the viscosity experienced by the bead and obtained using Stokes’ law γ = 6πηr, and τ_2_ = γ_2_/κ represents the characteristic time scale of the fluid. Two sets of simulations were performed to investigate the effect of cell size and shape, at different stages on the viscous drag and restoring stiffness. In the first set of simulations, the cytoplasm properties were fixed on their corresponding values of the first cell stage, and cell shape was assumed to be round, and only the cell and spindle size were modified. In a second set, the cytoplasm properties were fixed on their corresponding values of the first cell stage, and cell shapes, size and spindles size were changed according to experiments (Fig. 2 and Fig. S2). The shape of the cell boundary in the first division stage was a sphere with the radius R. To simplify cell shape modifications during cell division, we assumed that in the second and fourth division stages, the cell shapes were equivalent to half and one-fourth of a sphere with the same radius as the first cell stage. To simulate eggs in the microchambers, the width and length of the rectangular cube confinements were inserted from the cell shape in the experiments and the height was calculated using the volume of the spherical cells. Moreover, a set of simulations was done to inspect the consequence of the cross-section on the viscoelastic properties in the rectangular chambers (Figs. S6F-G). In this series, the volume of the chambers was the same as a spherical cell at the first division stage. The length of the chambers was constant and only the cross-section was changed from a square to a long rectangle. The spindle was considered as a rigid elastic (prolate) ellipsoid with long semi-axis a and short semi-axis b which was placed in the origin of the coordinate system in the geometrical center of the cells. The spindle dimensions were constant at each cell division stage in the simulations but were slightly reduced from the first to fourth stage, as in experiments. The cytoplasm was interacting with the pulled spindle by including viscoelastic flow and solid mechanics module in COMSOL through stress at the boundaries. A Heaviside step function was used to apply a constant force as a boundary load on half of the ellipsoid. The force direction was along the long axis of the ellipsoid for the translational displacement and perpendicular to the long axis at the initial time for the orthogonal displacement. The amplitude of loads in the time-dependent model was equal to the average magnetic forces in the experiments applied for 150 seconds and then released. The boundary conditions were no-slip and simulations were performed using a normal mesh size.

The viscoelastic parameters of simulation results were derived from the creep curve, as experimental curves, and fits of the Jeffreys model were performed by optimization method with a custom-written MATLAB script. Only the first frame of the pulling phase was used for the measurement of the vortex sizes in the simulations. The input parameters of the simulations are summarized in Table S1.

---

**Figure S1 (related to Figure 1).**
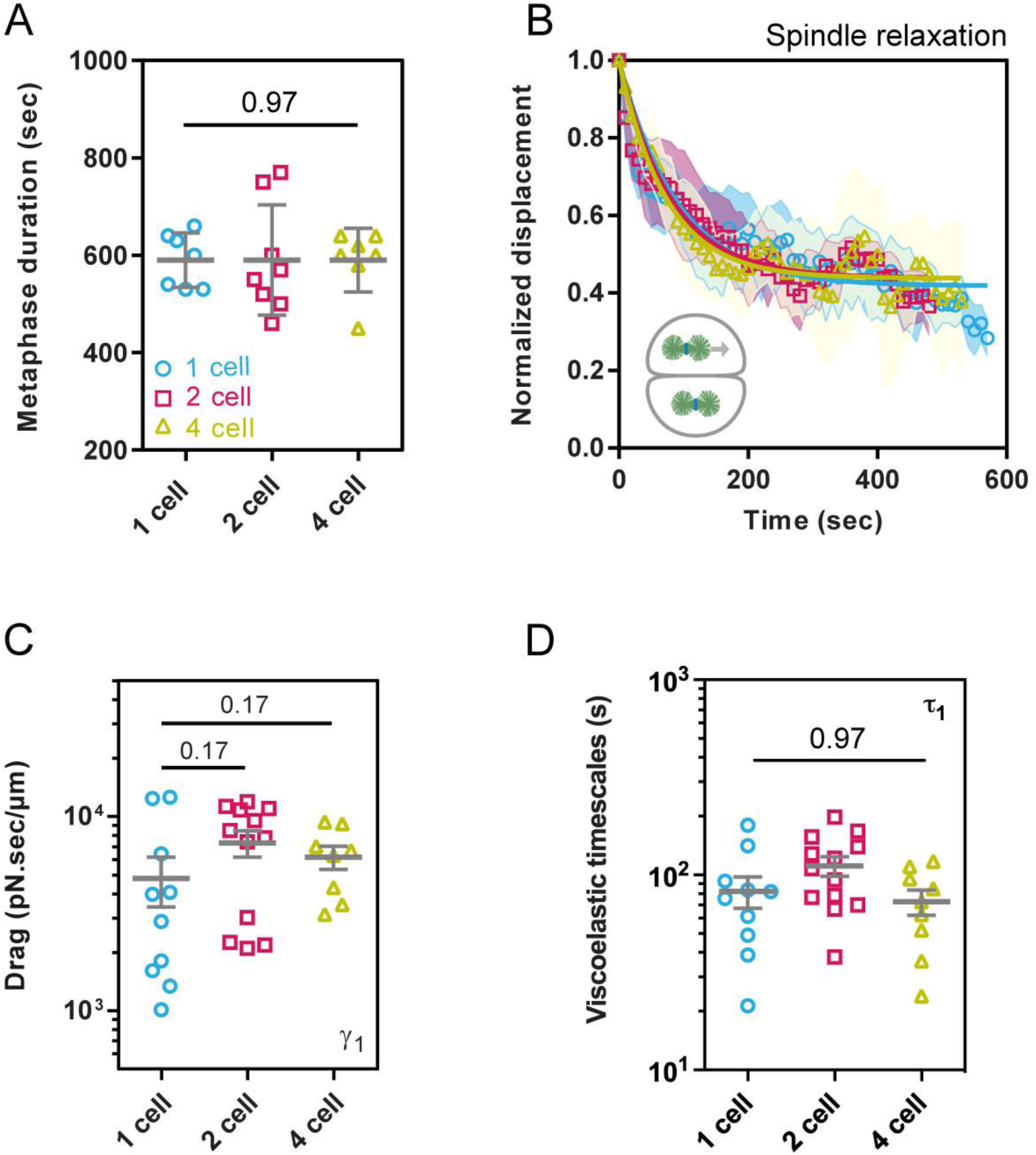
Dynamic of spindle recoiling back to the cell center upon force-induced displacement, and evolution of spindle drags and viscoelastic time-scales with cell stages. **(A)** Duration of metaphase in 1-cell, 2-cell and 4-cell embryos (n=7, 8 and 7 cells respectively). **(B)** Time evolution of the spindle displacement back to the cell center when the external magnetic force is released, normalized to that at the moment of force cessation, for different cell stages (n=10, 12 and 8 cells respectively). The bold lines correspond to fits of the data using relaxation equations of the Jeffreys’ viscoelastic model. Error bars are represented as shades in these curves and correspond to +/- SD/2. **(C)** Spindle drag coefficient moved parallel to the spindle axis in the indicated stages (n=10, 12 and 8 cells respectively), computed using fits to the Jeffreys’ models. **(D)** Viscoelastic time-scales of the Jeffreys’ model at different developmental stages. Error bars correspond to +/- SEM. Results were compared by using a two-tailed Mann– Whitney test. The corresponding P-values are reported on the graph. Scale bars, 20 μm. Error bars correspond to +/- SD unless otherwise indicated.

**Figure S2 (Related to Figure 2).**
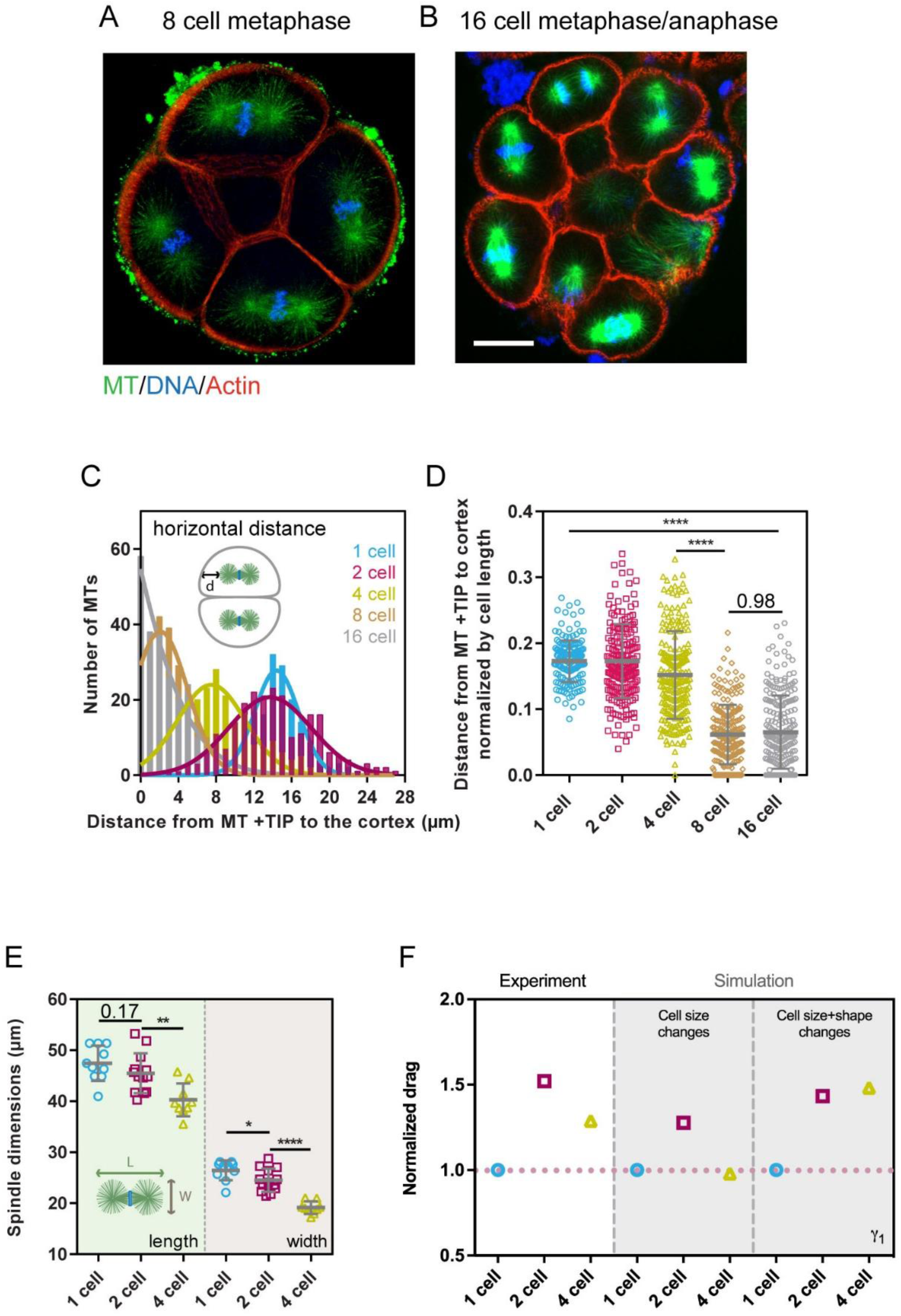
Comparison between experimental data and simulation of spindle restoring stiffness and drag. **(A-B)** Airy-scan confocal images of sea urchin embryos at 8-cell stage (A) and at 16-cell stage (B) in metaphase fixed and stained for Microtubules (MTs), DNA and F-actin. **(C)** Quantification of the distance from MT+ TIPs to the actin cortex in 1-cell, 2-cell, 4-cell, 8-cell and 16-cell embryos at metaphase (n= 168 MTs from 4 embryos, 229 MTs from 4 embryos, 220 MTs from 4 embryos, 223 MTs from 1 embryo and 230 MTs from 2 embryos respectively). **(D)** Quantification of the distance from MT+ TIPs to the actin cortex normalized by cell length for the same cells and conditions as in C. **(E)** Metaphase spindle length and width measured as indicated in the inset in F for different cell stages (n= 10, 13 and 9 cells respectively). **(F)** Spindle drag coefficient normalized by the 1-cell value compared between data from experiment and different sets of simulation at different developmental stages. Results were compared by using a two-tailed Mann–Whitney test. The corresponding P-values are indicated in the plot, or indicated as *, P<0.05, **, P<0.01, ***, P < 0.001, ****, P < 0.0001. Scale bars, 20 μm.

**Figure S3 (related to Figure 3).**
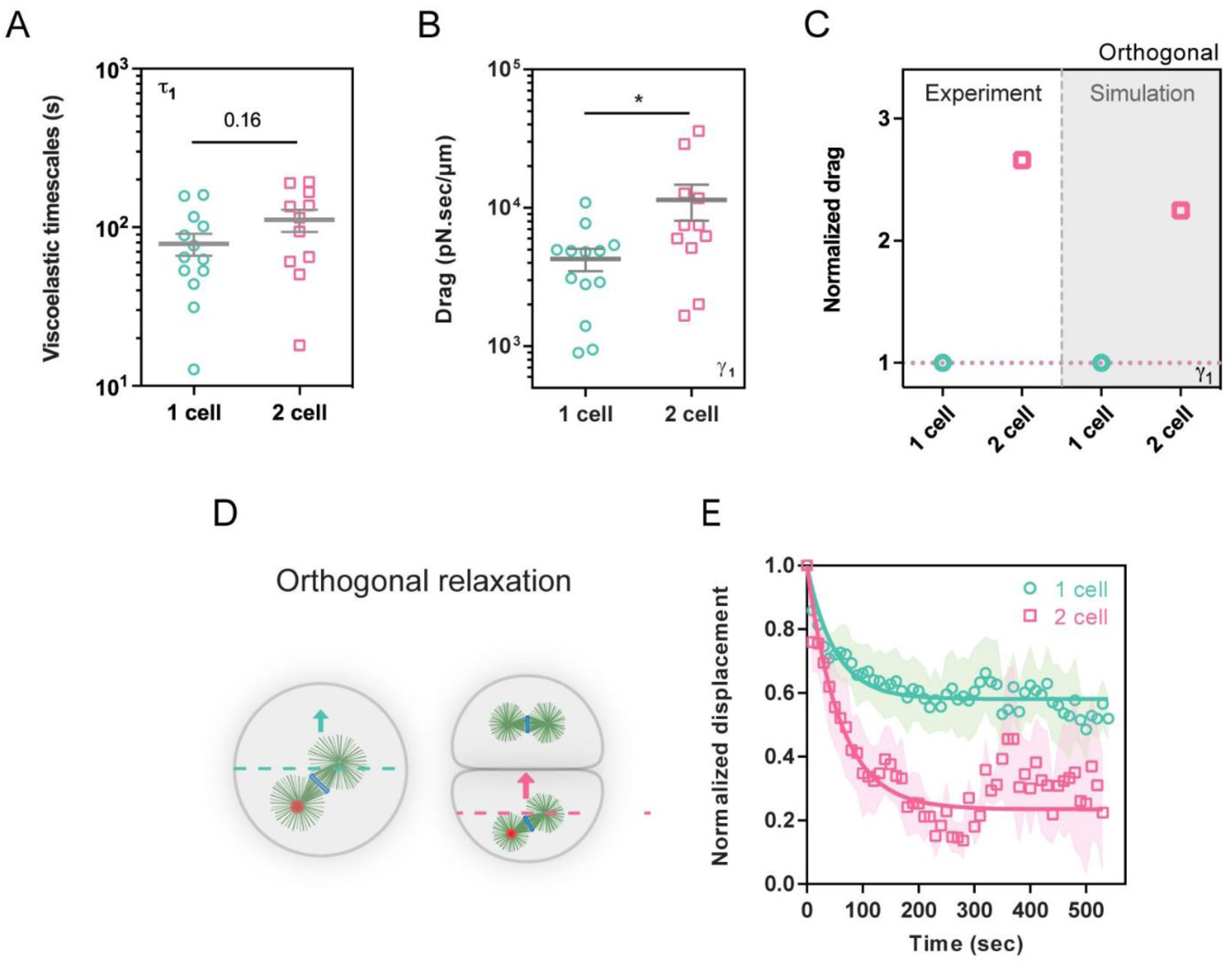
Dynamic of spindle recoiling back to the cell center upon force-induced displacement, and evolution of spindle drags and viscoelastic time-scales with cell stages. **(A)** Viscoelastic time-scales of the Jeffreys’ model at different developmental stages for spindles pulled orthogonal to their initial axis. Error bars correspond to +/- SEM. **(B)** Spindle drag coefficient for spindles moved orthogonal to the spindle axis in the indicated stages (n= 13, and 11 cells respectively), computed using fits to the Jeffreys’ models. **(C)** Spindle drag coefficient normalized by the 1-cell value compared between data from experiment and simulation at different developmental stages. **(D)** Scheme representing spindle relaxation upon force application orthogonal to the spindle axis. **(E)** Time evolution of the spindle displacement back to the cell center when the external magnetic force is released, normalized to that at the moment of force cessation, for different cell stages (n= 14 and 9 cells respectively). The bold lines correspond to fits of the data using relaxation equations of the Jeffreys’ viscoelastic model. Error bars are represented as shades in these curves and correspond to +/- SD/2. Results were compared by using a two-tailed Mann–Whitney test. Results were compared by using a two-tailed Mann–Whitney test. The corresponding P-values are indicated in the plot, or indicated as, *, P<0.05. Error bars correspond to +/- SD unless otherwise indicated.

**Figure S4 (related to Figure 4).**
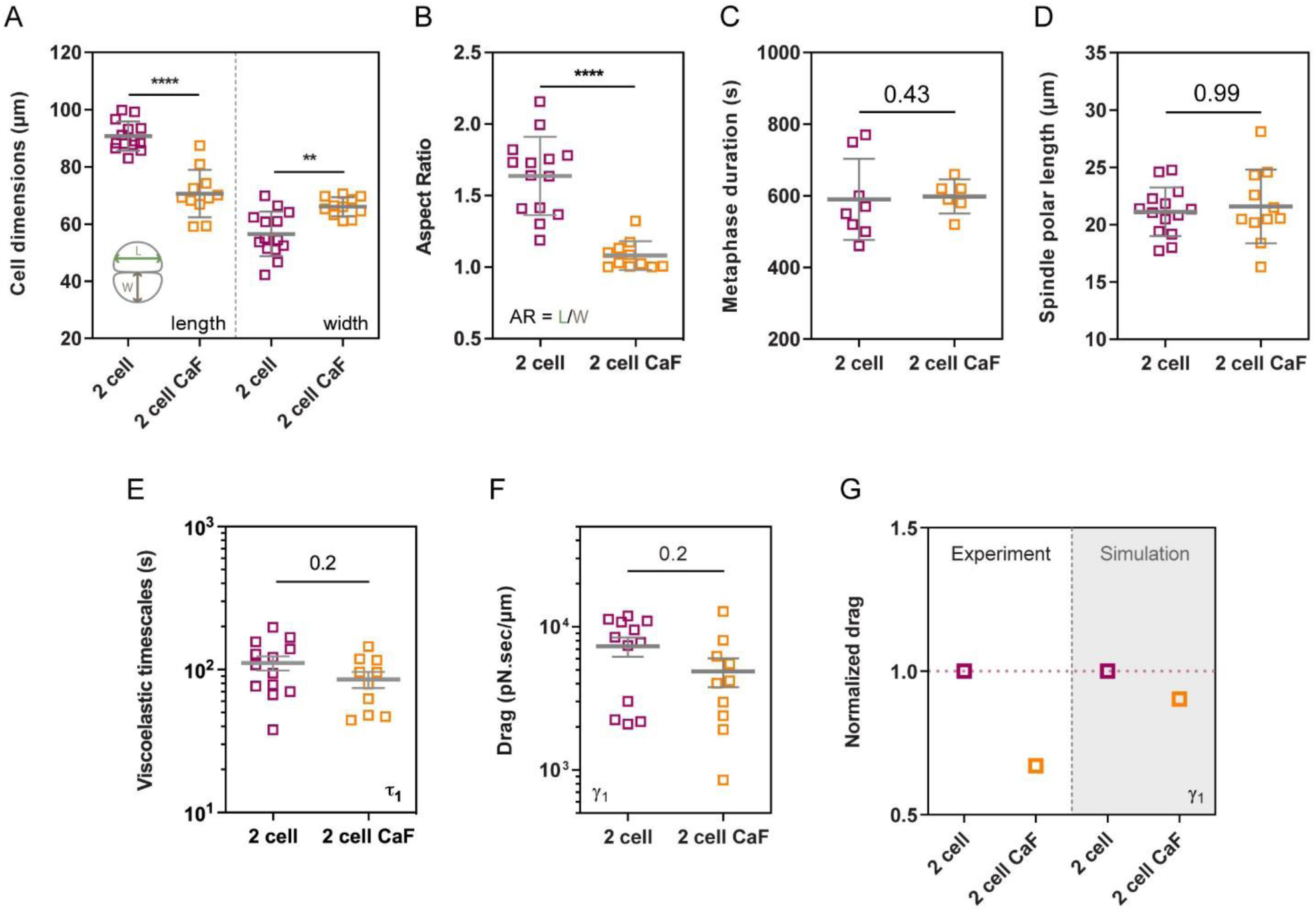
CaFSW treatment affects cell shapes and spindle drags in the cytoplasm. **(A)** Quantification of cell length and width for control cells vs cells treated with CaFSW at the 2-cell stage (n=14 and 11 cells respectively). **(B)** Metaphase duration for control cells vs cells treated with CaFSW at the 2-cell stage (n=8 and 6 cells respectively). **(C)** Spindle polar length for control cells vs cells treated with CaFSW at the 2-cell stage (n=14 and 11 cells respectively). **(D)** Spindle drag coefficient measured from metaphase spindle pulls for the indicated conditions (n=12 for and 10 cells respectively). Error bars correspond to +/- SEM. **(E)** Viscoelastic time-scales measured from metaphase spindle pulls for the indicated conditions (n=13 for and 10 cells respectively). Error bars correspond to +/- SEM. **(F)** Drag coefficient normalized by the control 2-cell value in indicated conditions for experimental data vs simulations. Results were compared by using a two-tailed Mann–Whitney test. The corresponding P-values are indicated in the plot or indicated as **, P < 0.01, ****, P < 0.0001. Error bars correspond to +/- SD unless otherwise indicated.

**Figure S5 (related to Figure 5).**
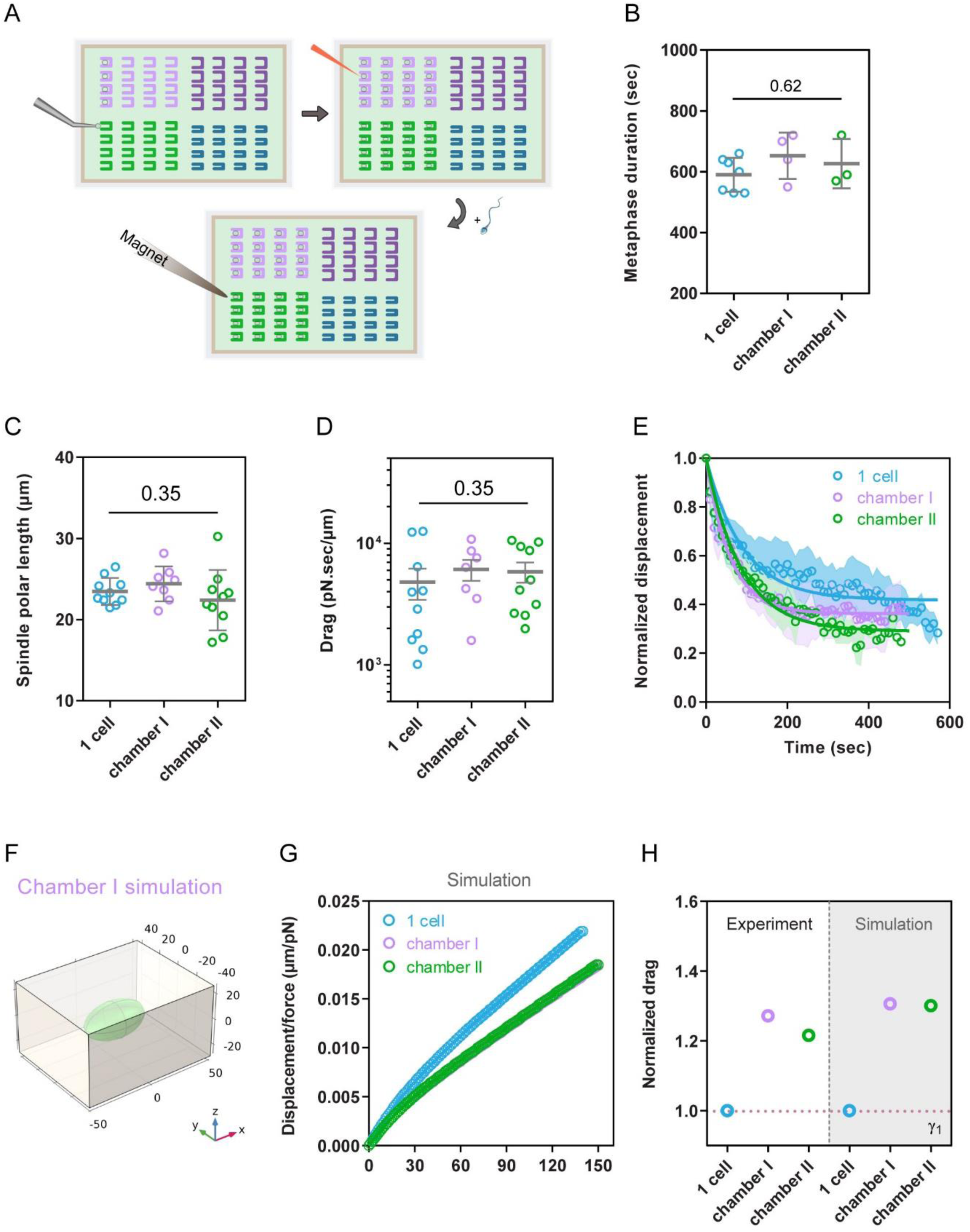
Impact of cell shape anisotropy on spindle movement and drags. **(A)** Scheme representing the process of spindle pulls in cells shaped in resin-made U chambers. First unfertilized eggs, are inserted in different protamine coated chambers using a glass pipette (top left), then magnetic beads are injected into eggs and eggs are fertilized (top right) lastly, forces are applied to beads bound to mitotic spindles with a magnetic tip (bottom). **(B)** Metaphase duration for cells at 1-cell stage in controls vs cells deformed in chambers of different aspect ratios (n=7, 4 and 3 cells respectively). **(C)** Spindle polar length for control cells vs cells shaped in chambers at 1-cell stage (n=16, 8 and 10 cells respectively). **(D)** Viscous drag measured from metaphase spindle pulls for control cells vs cells shaped in chambers at 1-cell stage (n=10, 8 and 10 cells respectively). Error bars correspond to +/- SEM. **(E)** Time evolution of normalized spindle recoil for control cells vs cells shaped in chambers at 1-cell stage (n=10, 8 and 10 cells respectively). Error bars are represented as shades in these curves and correspond to +/- SD/2. **(F)** Scheme representing the 3D simulation of spindle pulls in an egg confined in Chamber I, considering the cell shape as a rectangular cube. **(G)** Time evolution of the spindle displacement normalized by force in the indicated conditions predicted from simulations. **(H)** Viscous drag normalized by the 1-cell value compared between data from experiment and simulations. The red dotted line corresponds to value 1. Results were compared by using a two-tailed Mann–Whitney test. The corresponding P-values are indicated in the plot. Error bars correspond to +/- SD unless otherwise indicated.

**Figure S6 (related to Figure 5).**
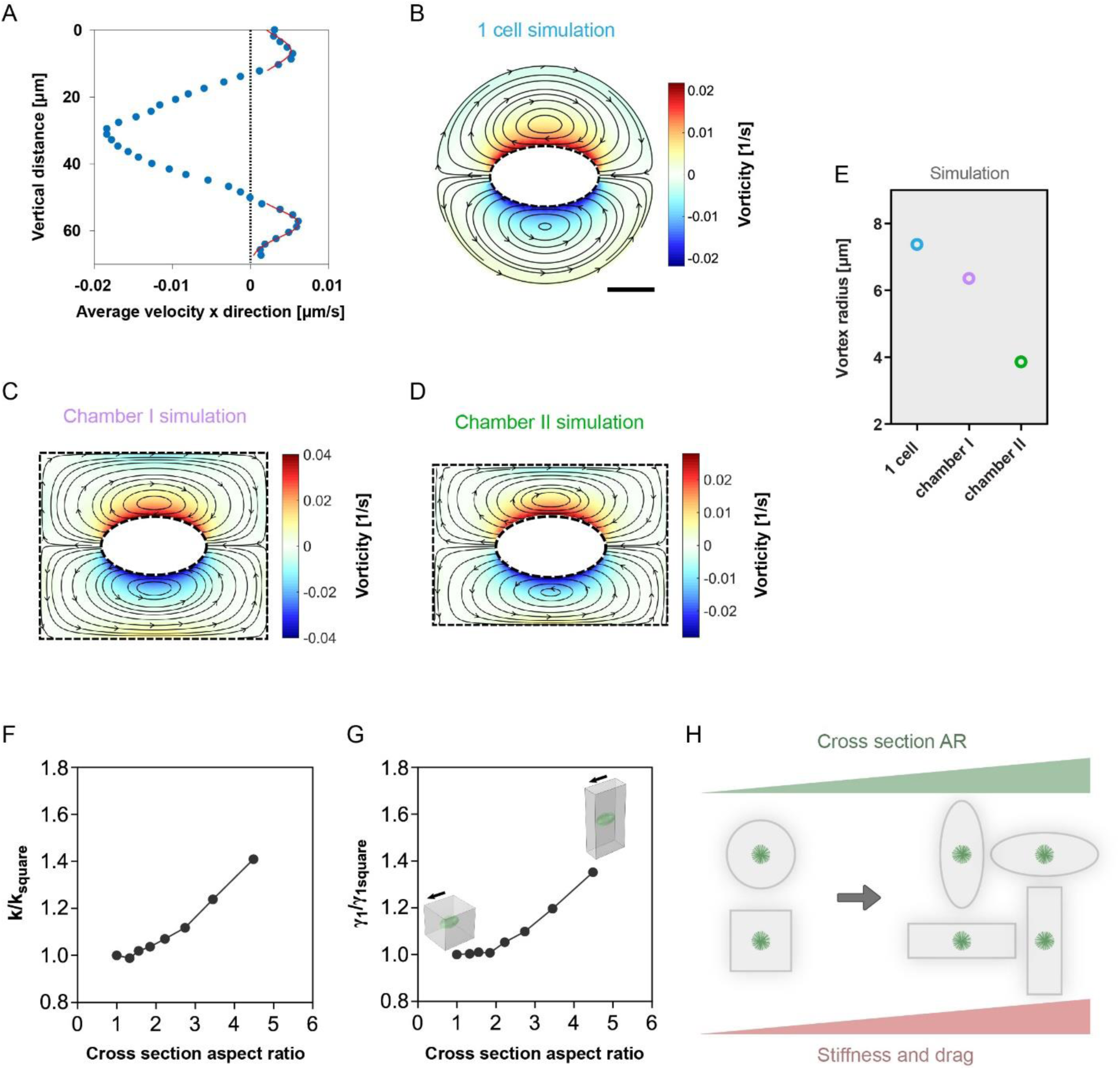
Analysis of cytoplasmic flows and associated simulations during spindle movement in different cell shapes. **(A)** The spatial average of the velocity component along the pulling direction (x), plotted as a function of the vertical distance (y). Red curves are Gaussian fits to compute the size of vortices. **(B-D)** Numerical cytoplasm streamlines during the first frame of spindle pulls in control cells (B), cells confined in chamber I (C), and cell confined in chamber II (D). Streamlines are superimposed onto a color map of local flow vorticities, ω. **(E)** Flow vortex sizes from simulations for cells confined in indicated rectangular cubes and unconfined cells. **(F-G)** Numerical prediction of the evolution of restoring stiffness and drags, as a function of the cross-sectional aspect ratio of the cell shape. Values are normalized to that of a square cross-section. **(H)** Scheme representing the model for how cell geometry may impact cytoplasm viscoelastic resistance on spindle motions. The higher the aspect ratio of the cell cross-section, the greater the elastic and viscous resistance to spindle movement. Scale bars, 20 μm.

**Table S1.**
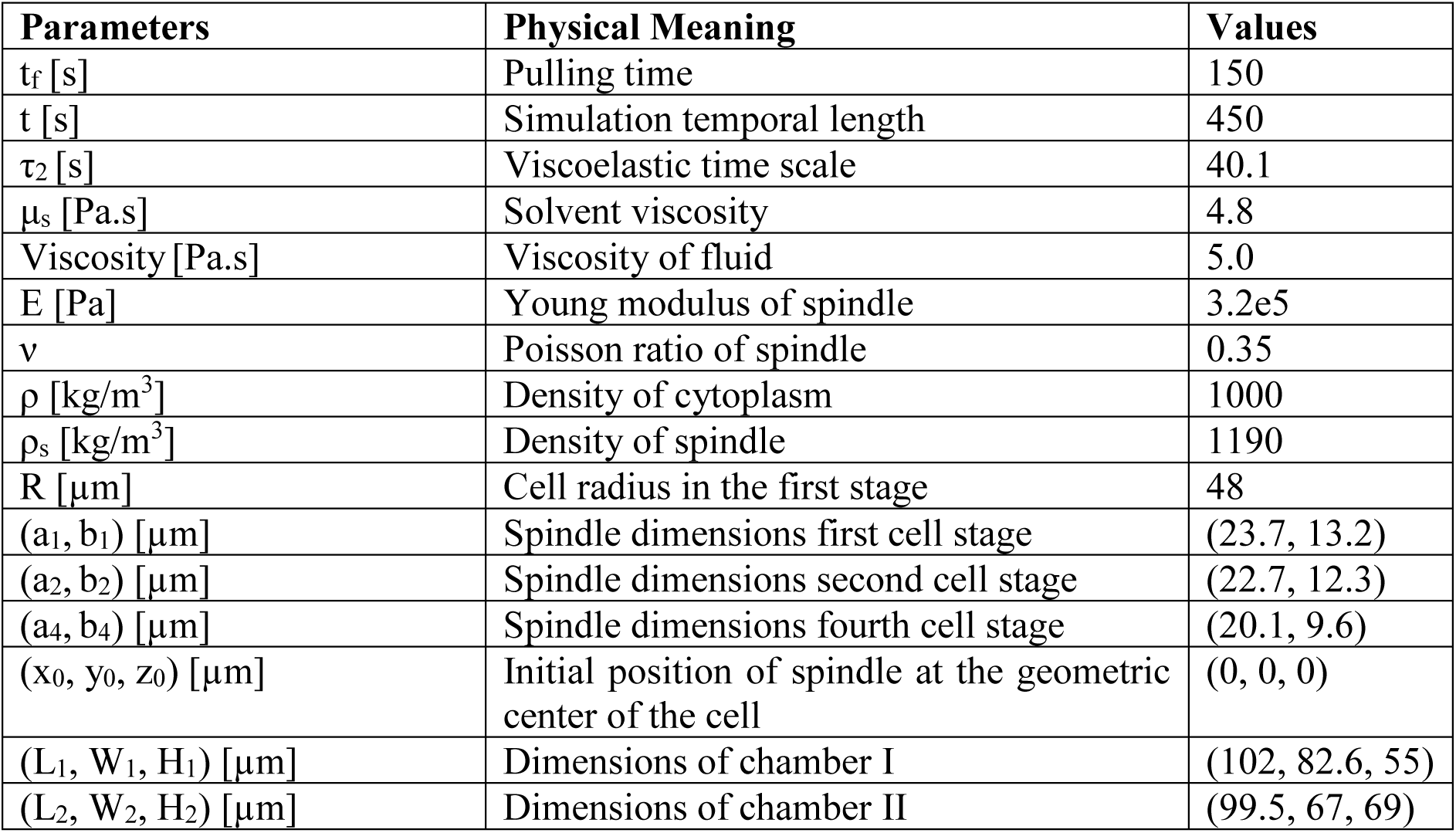
Parameters used in the 3D simulation of the spindle at different cell stages (fluid viscoelastic properties are for the first cell stage)

## Supplemental Movie legends

**<u>Movie S1</u>. Early embryo development with blastomeres of reduced size and different shapes.** Time-lapse of two developing embryos with DNA labelled with Hoechst from 1-cell stage to 16-cell stage. Time is in hr:min and scale bars are 20 μm.

**<u>Movie S2</u>. Metaphase spindles pulled parallel to their long axis with magnetic tweezers at different cell stages.** Time-lapse of metaphase spindles pulled with magnetic forces along their long axis at different cell stages and let to recoil and divide. Beads are labelled in red and DNA in cyan. Time is in min:sec and scale bars are 20 μm.

**<u>Movie S3</u>. COMSOL simulation of spindles pulled parallel to their long axis at different cell stages.** Time-lapse from COMSOL simulation of spindles that were pulled at different cell stages for 150 seconds and then released. The observation plane is passing through the middle of the spindle placed in the geometric center of the cell. Time is in min:sec and Scale bars are 30 μm.

**<u>Movie S4</u>. Metaphase spindles pulled orthogonal to their long axis with magnetic tweezers at 1- and 2-cell stages.** Time-lapse of metaphase spindles pulled with magnetic forces orthogonal tp their long axis at 1-cell and 2-cell stages and let to recoil and divide. Beads are labelled in red and DNA in cyan. Time is in min:sec and scale bars are 20 μm.

**<u>Movie S5</u>. COMSOL simulation of spindles pulled orthogonal to their long axis at at 1- and 2-cell stages cell stages.** Time-lapse from COMSOL simulation of spindles that were pulled at 1- and 2-cell stages for 150 seconds and then released. The observation plane is passing through the middle of the spindle placed in the geometric center of the cell. Time is in min:sec and Scale bars are 30 μm.

**<u>Movie S6.</u> Metaphase spindle pulled in the CaFSW-treated round cell at 2-cell stage.** Time-lapse of a metaphase spindle in a 2-cell stage round cell after CaFSW treatment, pulled with magnetic tweezers and let to recoil and divide. Beads are labelled in red and DNA in cyan. Time is in min:sec and scale bars are 20 μm.

**<u>Movie S7.</u> Metaphase spindles pulled in cells deformed in microchambers of different shapes.** Time-lapse of metaphase spindles in zygotes confined in chambers of two sizes, pulled with magnetic tweezers and let to recoil and divide. Beads are labelled in red and DNA in cyan. Time is in min:sec and scale bars are 20 μm.

